# Pharmacokinetics and Efficacy of the FimH Inhibitor Sibofimloc in Preventing Necrotizing Enterocolitis in Preterm Piglets

**DOI:** 10.64898/2026.10.12.758546

**Authors:** Valeria Melendez Hebib, Brooklynn Earls, Barbara Stoll, Caitlin Vonderohe, Gregory Guthrie, Henry Joseph Haiser, Martin Paton, Suresh K. Balani, Matthias Hesse, Douglas Burrin

## Abstract

Necrotizing enterocolitis (NEC) is a disease in preterm infants associated with an increased abundance of Enterobacteriaceae. FimH, a mannose-binding adhesin on Enterobacteriaceae type 1 fimbriae, mediates mucosal attachment and TLR4-dependent inflammatory cascades. We evaluated the safety, pharmacokinetics, and efficacy of an oral FimH inhibitor (Sibofimloc) to reduce Enterobacteriaceae colonization and prevent NEC in preterm piglets. We performed a 9-d dose-escalation safety study (10, 25, 50 mg/kg BID) assessing absorption and tolerability and a 10-d randomized efficacy study. Piglets in the efficacy study received an infant formula diet with or without oral sibofimloc (40 mg/kg BID), or donor human milk (DHM) as a healthy control. NEC incidence, tissue histopathology, and gut microbiota composition via metagenomic whole-genome sequencing (WGS) were evaluated. Systemic exposure data reveal that sibofimloc was dose-dependently absorbed, while plasma levels decreased from postnatal day 2 to day 8, indicating reduced drug absorption with maturation of the intestinal barrier. Treatment with sibofimloc did not reduce NEC incidence (100% in sibofimloc in formula controls) or severity, whereas DHM-fed piglets experienced 0% NEC. Sibofimloc-treated animals exhibited significantly reduced crypt depths, demonstrating heightened mucosal injury and impaired regenerative capacity. WGS revealed that sibofimloc-treated animals maintained a gut luminal microbiota composition similar to untreated formula controls, dominated by *Clostridium perfringens* (*C. perfringens*). Selective targeting of FimH-expressing Enterobacteriaceae was insufficient to prevent NEC or alter the gut microbiota in this preterm piglet model, underscoring the complexity of NEC-associated microbial communities and suggesting that additional microbial targets and other colonization mechanisms may contribute to disease development.

## Introduction

Necrotizing enterocolitis (NEC) is the most common gastrointestinal emergency in preterm infants, characterized by intestinal inflammation, necrosis, and a high mortality rate. The pathogenesis of NEC is multifactorial, yet evidence suggests that diet and the gut microbiota are important risk factors for disease incidence and progression (1). An increased abundance of Enterobacteriaceae, a family of gram-negative bacteria, specifically members like *Klebsiella pneumoniae (K. pneumoniae)* (2, 3), has been associated with NEC, highlighting these bacteria as potential contributors to disease pathogenesis. A key mechanism that has linked the colonization of Enterobacteriaceae with intestinal inflammation is by activation of toll-like receptor 4 (TLR4) which recognizes lipopolysaccharide (LPS) on the surface of these bacteria. TLR4 signaling in the gastrointestinal (GI) tract during the vulnerable perinatal period of preterm infants can impair mucosal/epithelial healing and contribute to bacterial translocation and systemic inflammation (4, 5).

FimH is a mannose-binding adhesin protein located at the tip of type 1 fimbriae on the surface of Enterobacteriaceae, such as *K. pneumoniae* and *Escherichia coli (E. coli)*. FimH facilitates bacterial adhesion to enterocytes by binding to mannose residues on the surface of epithelial cells, resulting in colonization. The importance of FimH in bacterial pathogenicity is most evident in infection with uropathogenic *E. coli* (UPEC) (6, 7). UPEC utilizes its abundantly expressed FimH adhesins to anchor itself to the bladder epithelium, enabling colonization, proliferation and biofilm formation within the bladder (6, 7). In inflammatory bowel disease, particularly Crohn’s disease, adherent-invasive *E. coli* (AIEC) becomes abnormally predominant within the ileal mucosa causing inflammation in the epithelium. FimH has been demonstrated to be essential for *E. coli* adhesion and subsequent inflammation through its interaction with TLR4 independently from LPS (8). Inhibition of FimH using a small molecule inhibitor (sibofimloc) has been demonstrated to reduce the ability of Crohn’s-associated clinical isolates of *E. coli* to adhere and cause inflammation *in vitro* using intestinal explants (9). In Crohn’s disease patients, oral administration of sibofimloc for 13 days resulted in a reduction of pro-inflammatory cytokines and calprotectin in the stool, demonstrating the importance of this adhesin in inflammation induced by AIEC (9). Due to the established association between an increase in Enterobacteriaceae colonization in NEC, we hypothesized that oral administration of sibofimloc would decrease the prevalence of Enterobacteriacea within the gut microbiota and therefore decrease the incidence and severity of NEC. To test this hypothesis, we used the preterm piglet, a well-established and physiologically relevant model of NEC in preterm infants(10). This translational preclinical model allowed us to evaluate the suitability, safety, and efficacy of sibofimloc to prevent NEC in a preterm neonate. We first assessed the tolerance, absorption, metabolism, and distribution of sibofimloc in a pharmacokinetic study and also tested the efficacy of oral sibofimloc treatment to prevent NEC in preterm piglets.

## Results

### SIBOFIMLOC dose-escalation pharmacokinetics and NEC outcomes in preterm piglets

In Study 1, piglets treated with three escalating doses of either 10, 25, or 50 mg/kg, twice a day (bid) were monitored for a total of 9 days. Piglets received total parenteral nutrition and enteral feeding of formula according to Supplemental Table 1. A total of 6 piglets developed NEC and were euthanized during the study **(Figure 1a)**. There was no significant difference in NEC-free survival or NEC severity between doses of sibofimloc **(Figure 1b-1c)**. In Study 1, absorption of sibofimloc was assessed on days 2 and 8 of the experiments by measuring the amount of sibofimloc in plasma. Plasma levels of sibofimloc were highest in the piglets that received the 50 mg/kg dose, followed by piglets that received 25 mg/kg dose on day 2 of life **(Table 1)**. A similar, dose dependent pattern was observed on day 8; piglets that received 50mg/kg of sibofimloc exhibited the highest plasma concentration of sibofimloc, however, the plasma concentration of the drug in groups 25 mg/kg and 10 mg/kg was similar. Notably, plasma concentration of sibofimloc significantly decreased with age in piglets receiving 50 mg/kg as evidenced by comparisons of area under the curve (AUC) and maximum concentration in plasma (C_max_) within doses from day 2 to day 8 of life **(Table 1) (Figure 2a and 2b**), suggesting reduced absorption of the drug correlates with decreased gut permeability with age. Comparison of Cmax and AUC among dose groups demonstrated a significant dose effect on DOL 2, whereas no significant differences in plasma exposure were observed among doses on DOL 8 **(Table 1)**. Calculation of the peak fraction of dose in blood (%) assuming a circulating blood volume of 70 ml/kg, revealed systemic absorption of sibofimloc is on average less than 2% across doses, not significantly different by dose but displaying dependency on postnatal age. This is evidenced by significant decrease in systemic absorption (to less than 0.2%) from DOL 2 to 8 at the higher doses (groups 25 and 50 mg/kg) **(Figure 2c)**.

**Figure 1.**
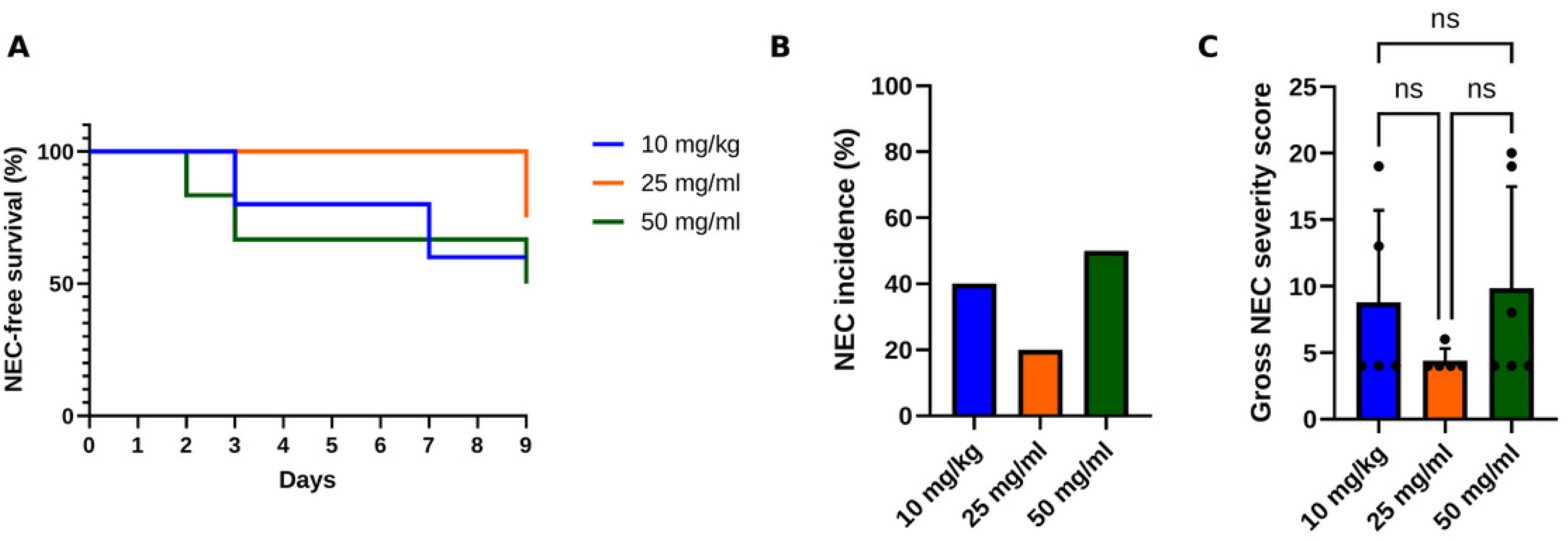
Pharmacokinetic dose-escalation study (Study 1): NEC incidence, survival, and severity are unaffected by sibofimloc dose. Preterm piglets received oral sibofimloc twice daily at 10, 25, or 50 mg/kg BID (n = 5-6 per group) and were monitored for necrotizing enterocolitis (NEC). (A) Cumulative NEC incidence. (B) Kaplan– Meier NEC-free survival. (C) Total gross NEC severity score; bars represent mean and each symbol represents one animal. NEC incidence was compared by Fisher’s exact test, NEC-free survival by log-rank (Mantel–Cox) test, and severity scores by one-way ANOVA with Tukey’s multiple-comparisons test. No significant differences were detected among doses (ns, not significant).

**Figure 2.**
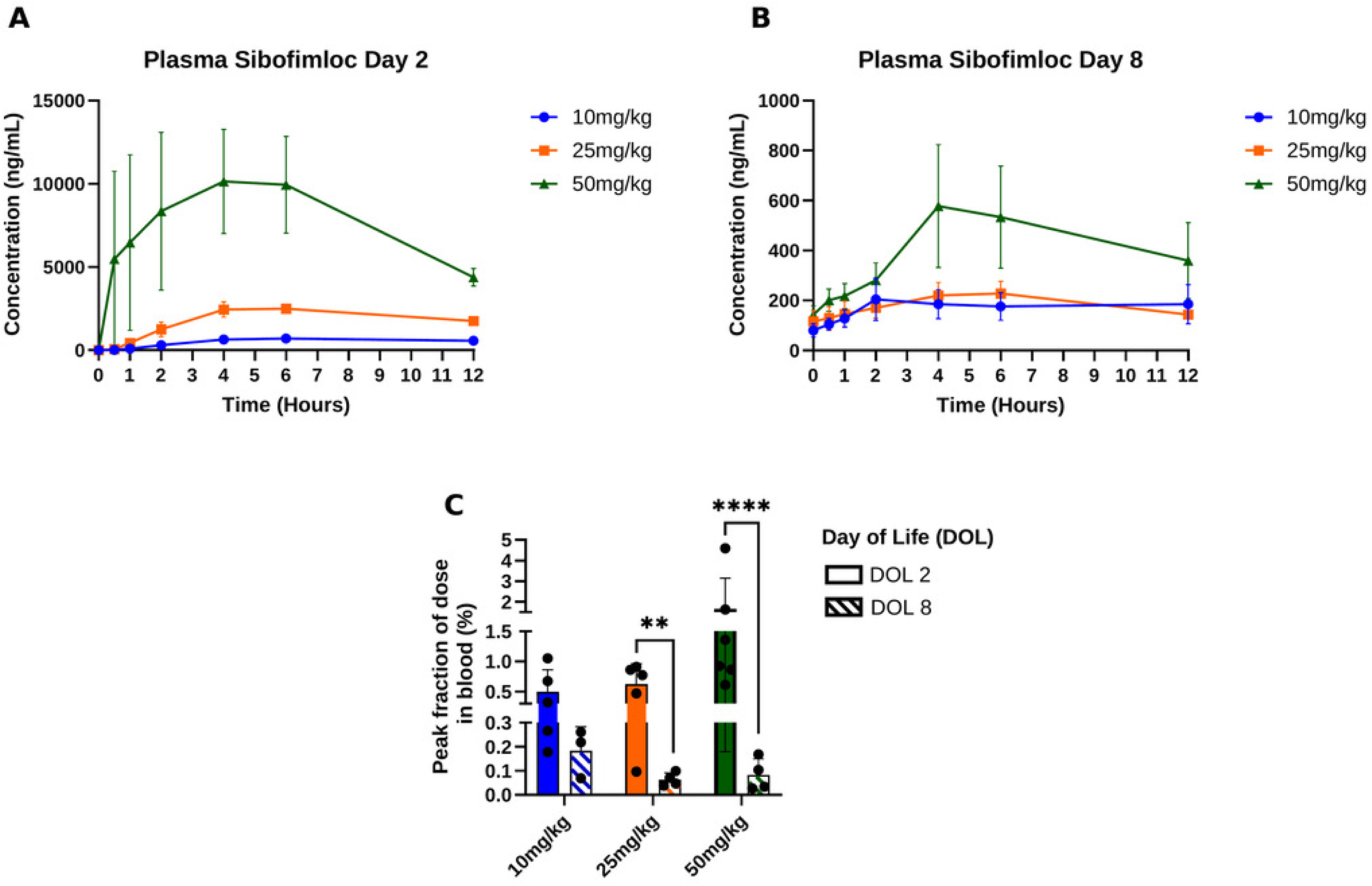
Systemic sibofimloc absorption is dose-dependent and declines with postnatal age. Plasma sibofimloc concentration–time profiles in Study 1 piglets dosed at 10, 25, or 50 mg/kg BID, sampled over 12 h post-dose. (A) Day of life (DOL) 2. (B) DOL 8. Points are group means; the y axis is plasma concentration (ng/mL) and the x axis is time post-dose (h). (C) Peak fraction of dose in blood (%) on DOL 2 and DOL 8. Percentages were log-transformed and compared using a two-way ANOVA with Šídák’s multiple comparisons test (✱✱, P < 0.01; ✱✱✱✱, P < 0.0001).

**Table 1.** Pharmacokinetics of SIBOFIMLOC doses in preterm piglets at DOL 2 and 8.

| Dose Level<br>(mg/kg) | DOL | Sex | N | T<sub>max</sub><sup>a</sup><br>(h) | C<sub>max</sub><br>(ng/mL) | P-value<br>(DOL 2 vs 8) | AUC<sub>0-12</sub><br>(h•ng/mL) | P-value<br>(DOL 2 vs 8) |
| --- | --- | --- | --- | --- | --- | --- | --- | --- |
| 10 | 2 | M+F | 5 | 6.0 (4.0-6.0) | 710±517<sup>a</sup> |  | 6390±4730<sup>a</sup> |  |
|  | 8 | M+F | 3 | 2.0 (2.0-12.0) | 261±143 | ns | 2110±1150 | ns |
| 25 | 2 | M+F | 4 | 5.0 (4.0-6.0) | 2690±708<sup>b</sup> |  | 22800±6430<sup>ab</sup> |  |
|  | 8 | M+F | 4 | 5.0 (4.0-6.0) | 228±97.8 | ns | 2240±942 | ns |
| 50 | 2 | M+F | 5 | 6.0 (4.0-6.0) | 7680±2920<sup>b</sup> |  | 62900±20700<sup>c</sup> |  |
|  | 8 | M+F | 4 | 4.0 (2.0-6.0) | 595±472 | 0.0095 | 5090±3710 | 0.0095 |

<sup>a</sup>Data are presented as mean ± SD unless otherwise indicated. Tmax values are presented as median (range).
Abbreviations: AUC<sub>0-12</sub>, area under the plasma concentration-time curve from 0 to 12 h; C<sub>max</sub>, maximum observed plasma concentration; T<sub>max</sub>, time to maximum plasma concentration; DOL, day of life.
P values represent comparisons between DOL 2 and DOL 8 within each dose group using the Mann–Whitney U test.
Means with different superscripts within a column are different ( $P \geq 0.05$ ). ns, not significant ( $P \geq 0.05$ ).

### Sibofimloc treatment does not reduce NEC incidence or severity in formula-fed piglets

In Study 2, twice-daily, oral treatment with 40 mg/kg of sibofimloc failed to reduce NEC incidence and severity in formula-fed piglets **(Figure 3a-3b)**. Sibofimloc administration yielded a NEC incidence of 100%, which is not significantly different than in formula-fed control (CTL) piglets (75%). On the other hand, NEC incidence in DHM-fed piglets (0%) was significantly lower than in both CTL and sibofimloc -treated piglets (P<0.05). Similarly, gross and histological NEC scores revealed no significant differences (P>0.05) in disease severity between sibofimloc treated- and CTL piglets, while DHM-fed piglets had significantly lower gross and histologic severity scores **(Figures 3c-3d)**. Additional analysis of H&E-stained sections of the distal ileum revealed no significant differences in villus height between treatments, however, NEC-afflicted piglets exhibited significantly lower villus height compared to healthy piglets regardless of treatment **(Figure 4a-4b)**. sibofimloc-treated piglets exhibited significantly lower crypt depths compared to both CTL piglets and DHM-fed piglets **(Figure 4c)**. Piglets afflicted by NEC had significantly smaller crypt depths compared to healthy piglets **(Figure 4d)**.

**Figure 3.**
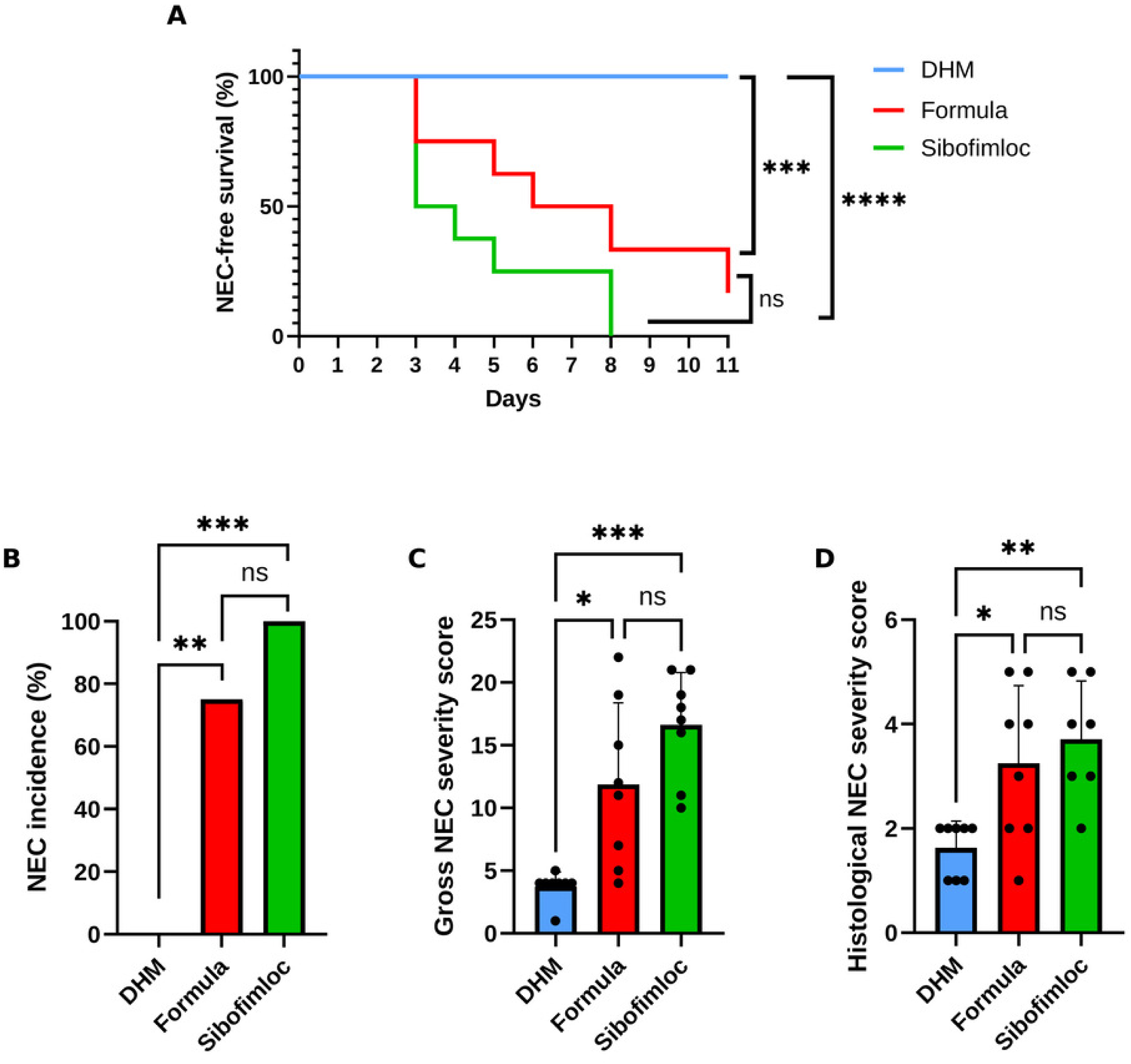
Sibofimloc does not reduce NEC incidence or severity in formula-fed piglets, whereas donor human milk is protective (Study 2). Preterm piglets were fed infant formula with vehicle (Formula/CTL), formula with oral SIBOFIMLOC (40 mg/kg BID; Sibofimloc), or donor human milk (DHM) (n = 7–8 per group). (A) Cumulative NEC incidence (100% Sibofimloc, 75% Formula, 0% DHM). (B) Kaplan–Meier NEC-free survival. (C) Total gross NEC severity score. (D) Distal ileum histological NEC severity score. In (C) and (D) each symbol is one animal and the bar is the group mean. NEC incidence was compared by Fisher’s exact test, survival by log-rank (Mantel–Cox) test, and severity scores by one-way ANOVA with Tukey’s multiple-comparisons test. (✱, P < 0.05; ✱✱, P < 0.01; ✱✱, P < 0.001; ns, not significant).

**Figure 4.**
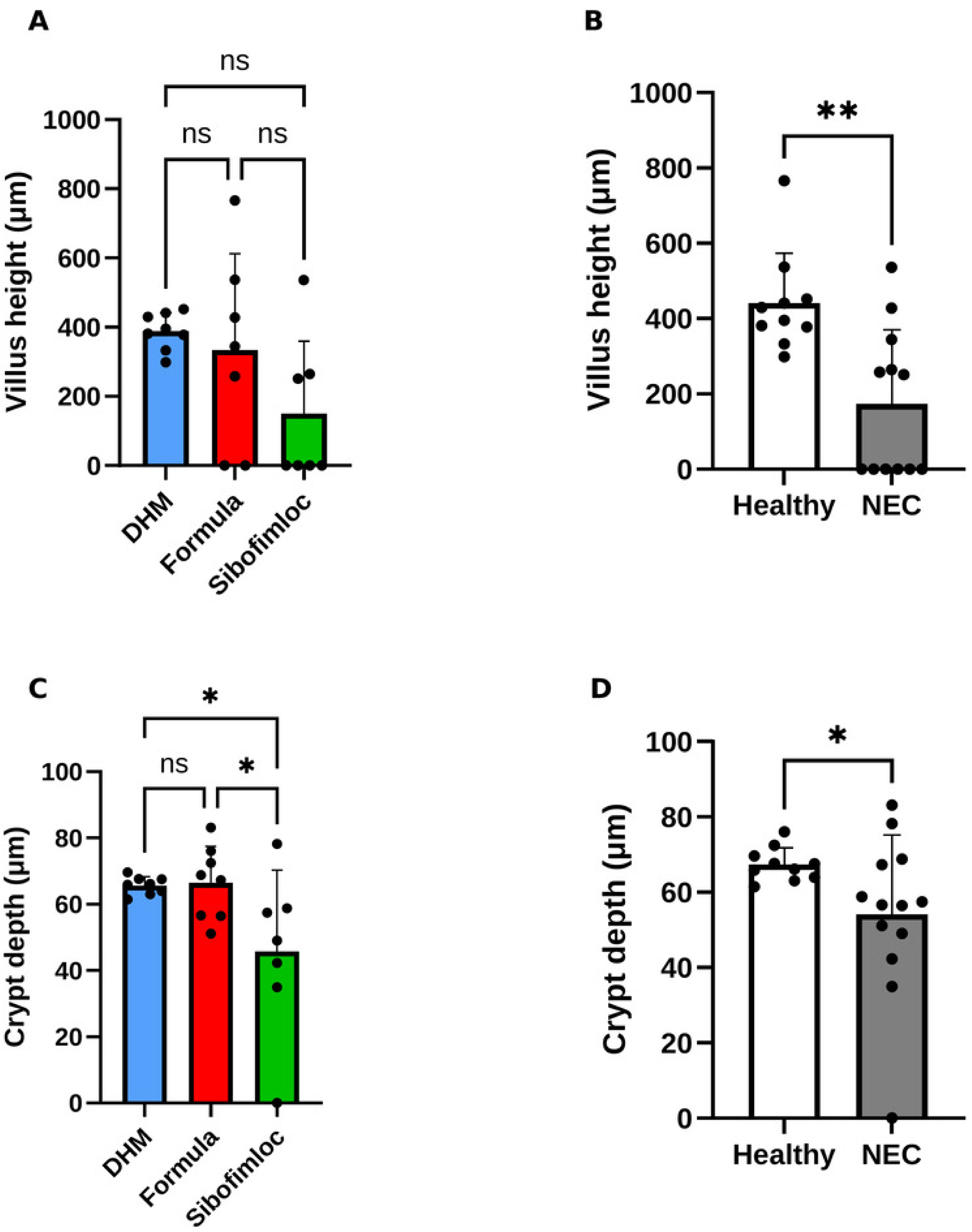
Sibofimloc reduces crypt depth, and NEC is associated with reduced villus height and crypt depth, in the distal ileum. Morphometry of H&E-stained distal ileum from Study 2 piglets. (A) Villus height by treatment (DHM, Formula, Sibofimloc). (B) Crypt depth by diagnosis; sibofimloc piglets had significantly lower crypt depth than both Formula and DHM piglets. (C) Villus height grouped by diagnosis (Healthy vs NEC). (D) Crypt depth grouped by diagnosis. Measurements are in micrometers (μm); each symbol is one animal and the bar is the group mean (treatment groups n = 7–8; Healthy n = 10, NEC n = 12–13). Treatment comparisons used one-way ANOVA with Tukey’s multiple-comparisons test; diagnosis comparisons used an unpaired t test with Welch’s correction (✱, P < 0.05; ✱✱, P < 0.01;ns, not significant).

### Treatment and diagnosis affect gut microbiota bacterial diversity

In Study 2, analysis of alpha diversity metrics on small intestinal and colonic contents revealed no significant differences in observed OTUs or Simpson diversity index between treatments nor between healthy and NEC-diagnosed piglets in the small intestine **(Figure 5a, 5b, 5e, 5f)**. On the other hand, in the colon, DHM-fed piglets exhibited a significantly higher observed OTUs compared to SIBOFIMLOC-treated piglets, whereas there was no significant difference between sibofimloc and CTL piglets **(Figure 5c-5d)**. Similarly in the colon, healthy piglets exhibited significantly higher observed OTUs and Simpson diversity index compared to piglets afflicted by NEC **(Figure 5g-5h)**.

**Figure 5.**
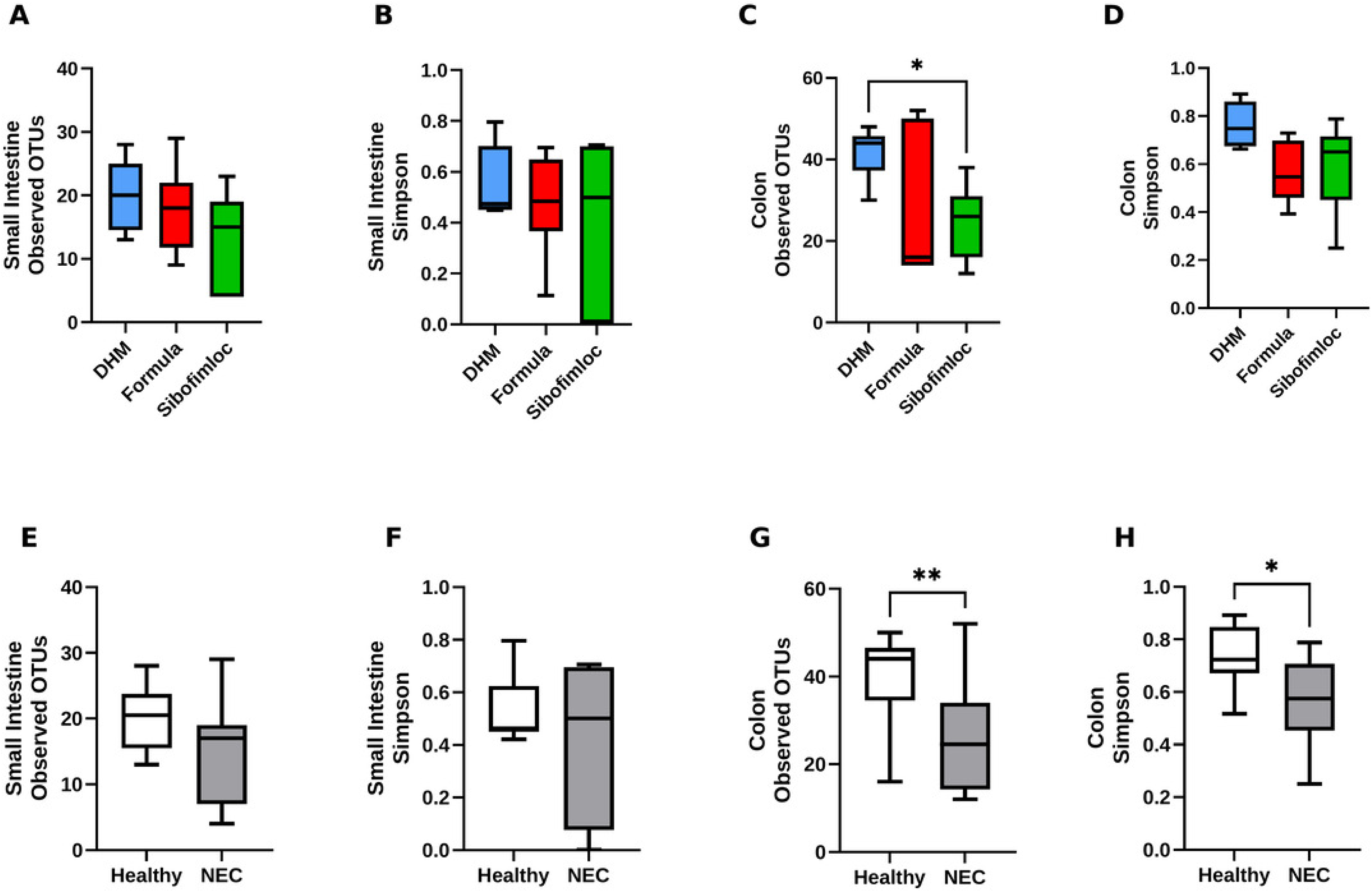
Gut microbial alpha diversity differs by diet and diagnosis in the colon but not the small intestine. Alpha diversity of luminal contents in Study 2, by treatment (DHM, Formula, Sibofimloc) and by diagnosis (Healthy vs NEC). (A) Observed OTUs, small intestine, by treatment. (B) Simpson index, small intestine, by treatment. (C) Observed OTUs, colon, by treatment; (D) Simpson index, colon, by treatment. (E) Observed OTUs, small intestine, by diagnosis. (F) Simpson index, small intestine, by diagnosis. (G) Observed OTUs, colon, by diagnosis. (H) Simpson index, colon, by diagnosis. Each symbol is one animal and the line is the group mean. Comparisons among treatments used the Kruskal–Wallis test with pairwise Wilcoxon rank-sum tests; Healthy-versus-NEC comparisons used the Wilcoxon rank-sum test (✱, P < 0.05).

### Treatment with SIBOFIMLOC did not alter the gut microbiota compared to untreated formula-fed piglets

Whole genome sequencing of small intestinal and colonic contents revealed that the gut microbiota composition of sibofimloc-treated piglets is similar to that of untreated CTL piglets **(Figure 6a-6b)**. For both sibofimloc-treated piglets and CTL piglets, *Clostridium perfringens (C. perfringens)* was the most abundant species within the gut microbiota **(Figure 6a, 6b, 7a)**. In the small intestine, the relative abundance of most gram-negative bacteria was suppressed by formula feeding and sibofimloc treatment, alike, highlighted by the significantly lower abundance of *Klebsiella pneumoniae (K. pneumoniae)*, *Klebsiella variicola (K. variicola)*, *Klebsiella quasipneumoniae (K. quasipneumoniae)*, and *Escherichia coli (E. coli)* compared to DHM-fed piglets **(Figure 7b-7e)**. DHM feeding also significantly increased gram-positive *E. faecalis* **(Figure 7f)**. In contrast, the colonic microbiota composition exclusively exhibited a significant difference of two bacterial species between NEC-positive and negative treatment groups. While sibofimloc-treated- and CTL piglets had a higher abundance of *C. perfringen*s in the colon, DHM-fed piglets were instead colonized by clostridium species, *Clostridium clostridioforme*, now known as *Enterocloster clostridioformis*, which was as abundant as >50% of the bacterial population in two piglets **(Figure 8b)**.

**Figure 6.**
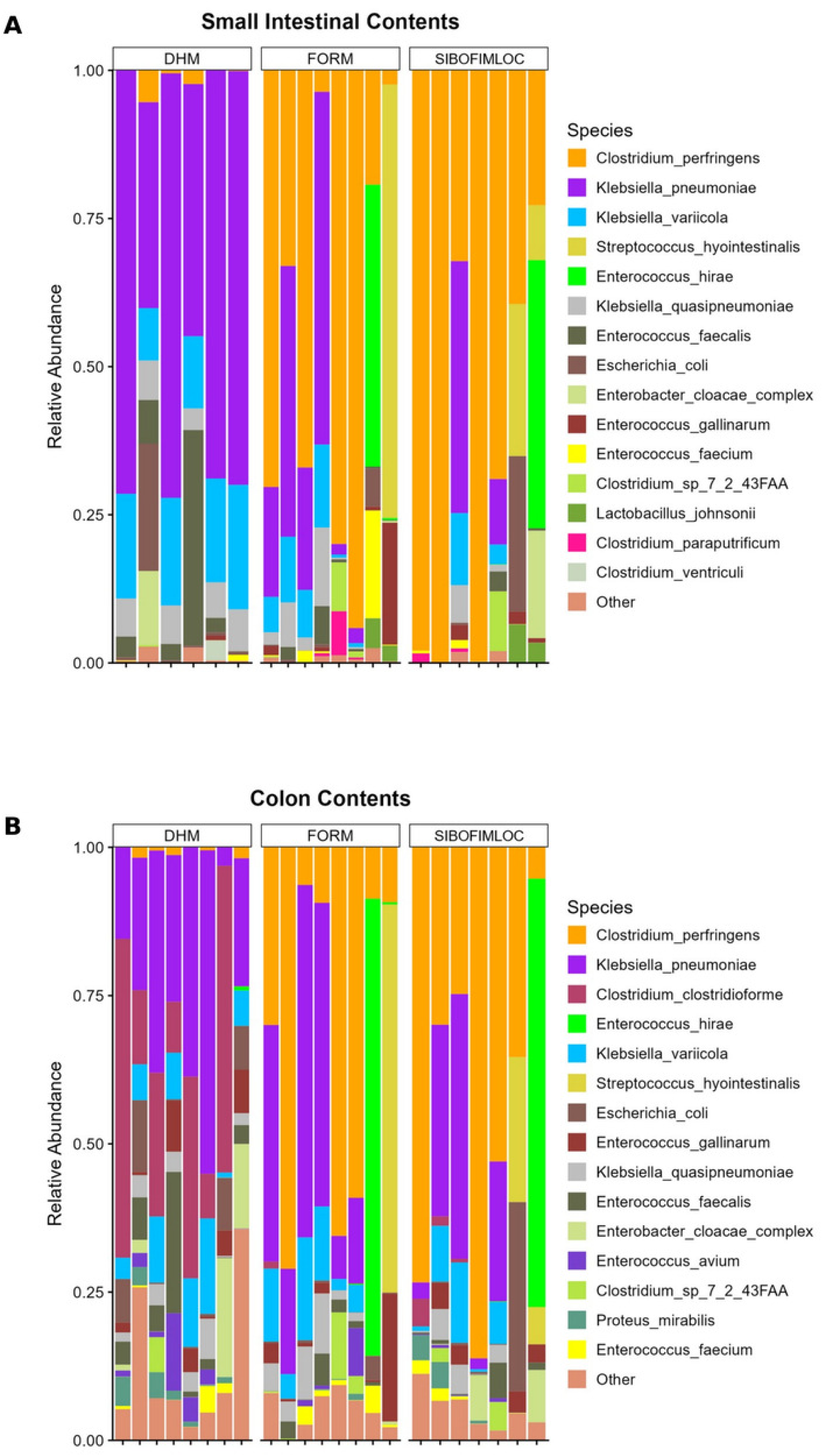
Sibofimloc does not alter overall gut microbiota composition relative to formula-fed controls. Relative abundance of bacterial species (metagenomic whole-genome shotgun sequencing) in luminal contents of Study 2 piglets by treatment (DHM, Formula [FORM], Sibofimloc). (A) Small intestinal contents. (B) Colon contents. Each stacked bar is one animal; colors denote species as indicated in the key, with low-abundance taxa collapsed into “Other”.

**Figure 7.**
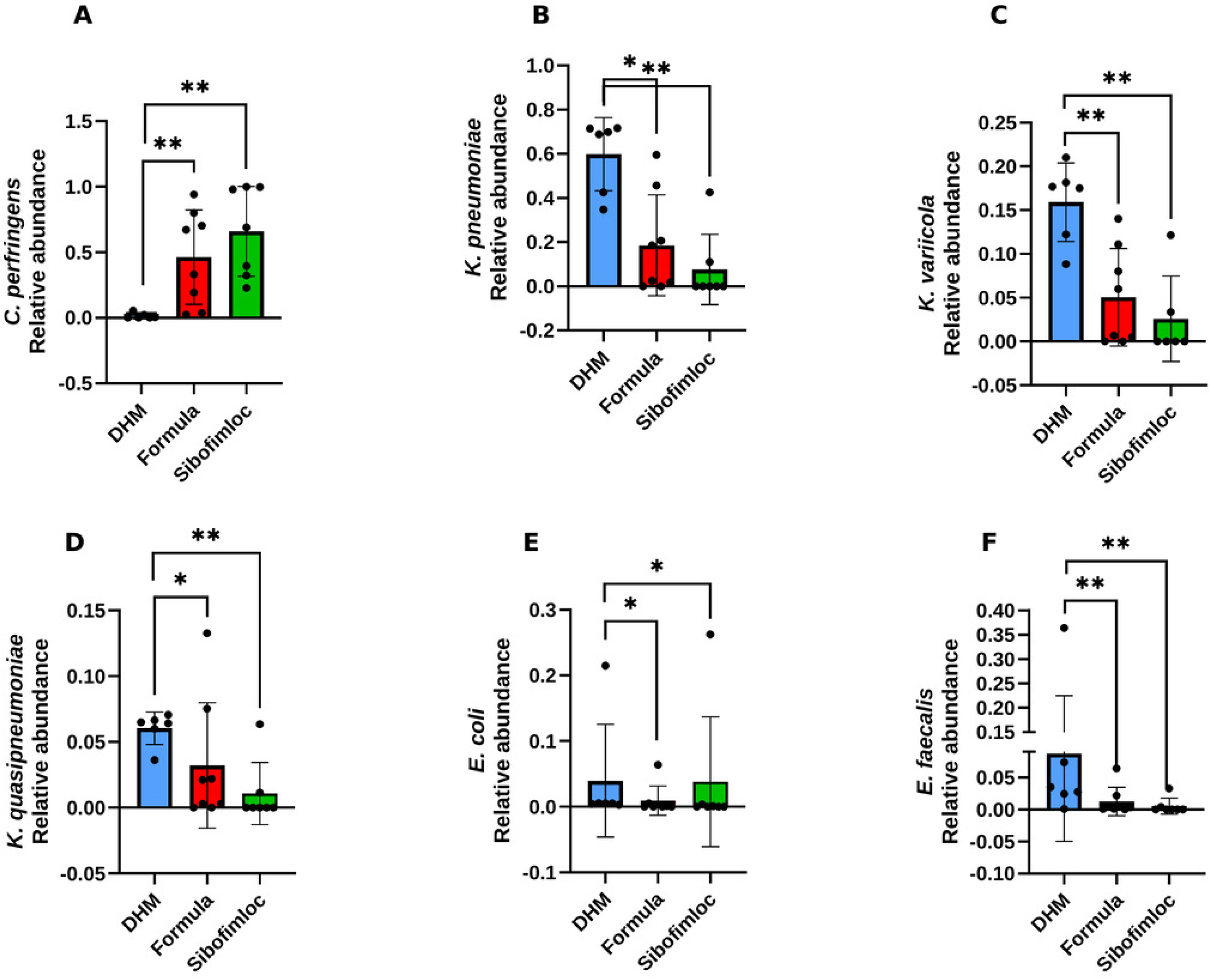
Formula feeding and sibofimloc suppress Enterobacteriaceae in the small intestine relative to donor human milk. Relative abundance of individual species in small intestinal contents of Study 2 piglets by treatment (DHM n = 6, Formula n = 8, Sibofimloc n = 6–7). (A) *Clostridium perfringens*. (B) *Klebsiella pneumoniae*. (C) *Klebsiella variicola*. (D) *Klebsiella quasipneumoniae*. (E) *Escherichia coli*. (F) *Enterococcus faecalis*. Each symbol is one animal and the bar is the group mean. Abundances were CLR-transformed and compared by Kruskal–Wallis test with pairwise Wilcoxon rank-sum tests; P values were adjusted by the Benjamini–Hochberg false-discovery-rate (FDR) method, with adjusted P < 0.05 considered significant (✱, P < 0.05; ✱✱, P < 0.01).

**Figure 8.**
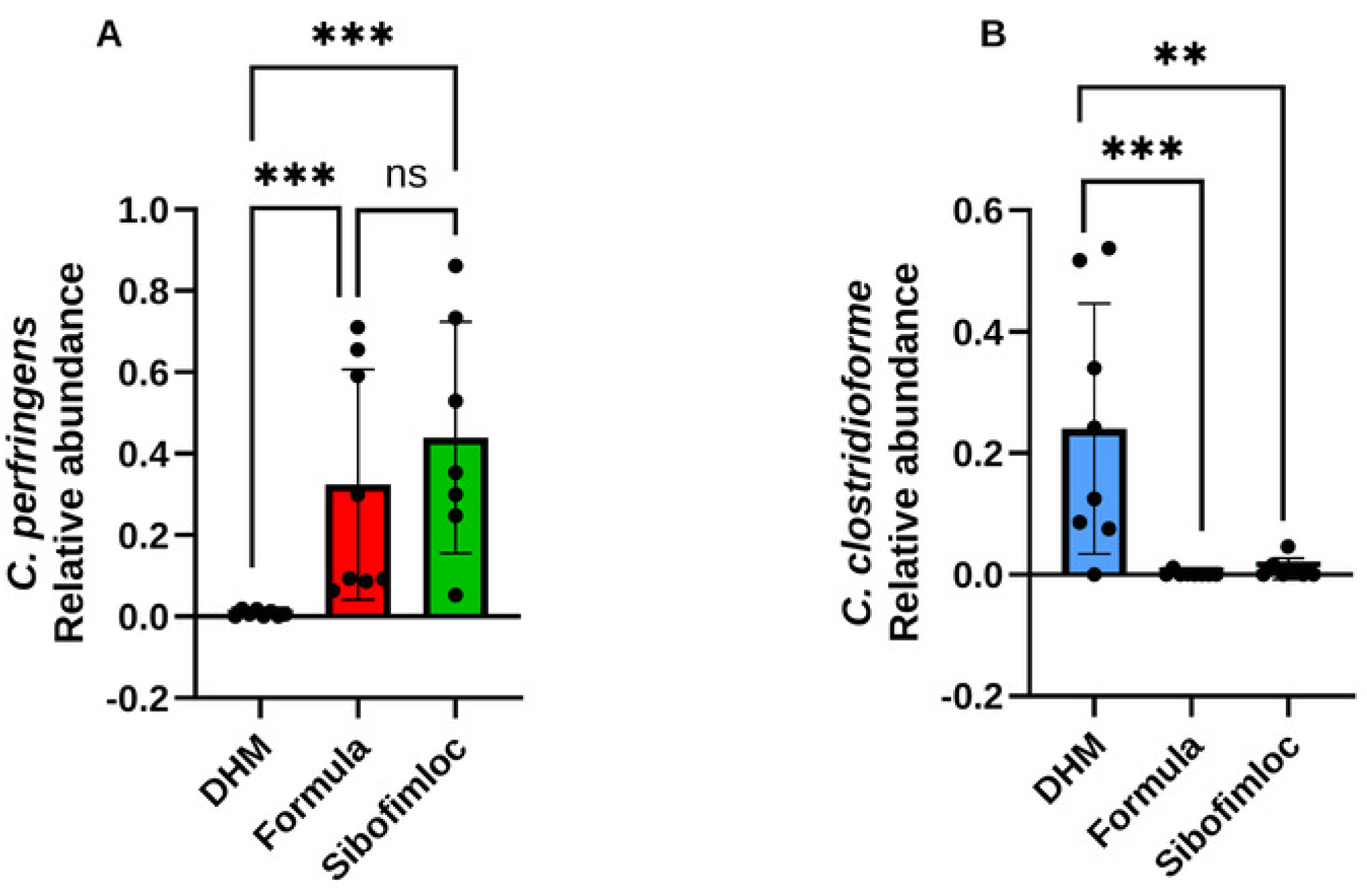
Colonic *Clostridium perfringens* dominates formula-fed and sibofimloc piglets, whereas donor human milk promotes *Enterocloster clostridioformis*. Relative abundance of individual species in colon contents of Study 2 piglets by treatment (DHM n = 8, Formula n = 8, Sibofimloc n = 7). (A) *Clostridium perfringens*. (B) *Clostridium clostridioforme* (reclassified as *Enterocloster clostridioformis*). Each symbol is one animal and the bar is the group mean. Abundances were CLR-transformed and compared by Kruskal–Wallis test with pairwise Wilcoxon rank-sum tests and Benjamini–Hochberg FDR correction (✱✱, P < 0.01; ✱✱, P < 0.001; ns, not significant).

### Clostridium perfringens abundance is associated with NEC

Analysis of gut microbiota composition by diagnosis (Healthy vs NEC), evidenced that *C. perfringens* made up the majority of the bacterial community in most piglets diagnosed with NEC, whereas healthy piglets exhibited more species diversity. *C. perfringens* abundance significantly increases in both the small intestine and the colon of piglets with NEC compared to healthy piglets **(Figure 9a, 9b, 10a, 11a)**. Piglets that did not develop NEC had a significantly higher abundance of *C. clostridioforme* and modest increases in the abundance of *K. pneumoniae, K variicola, K. quasipneumoniae,* and *E. faecalis* **(Figure 10b-10e)**. Abundance of *E. coli* did not notably differ between diagnoses **(Figure 10f)**.

**Figure 9.**
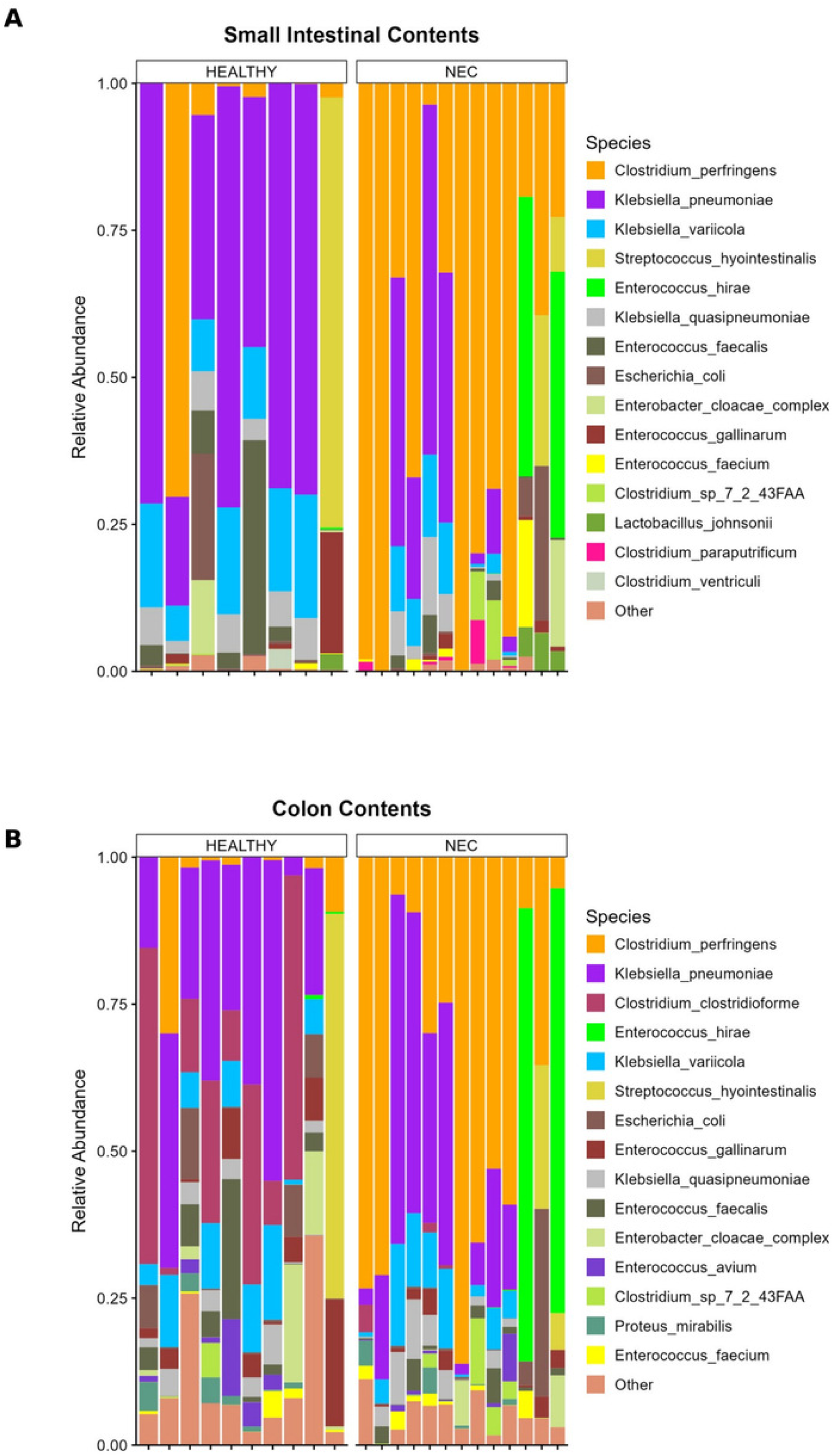
NEC is associated with a low-diversity microbiota dominated by *Clostridium perfringens*. Relative abundance of bacterial species (metagenomic whole-genome shotgun sequencing) in luminal contents of Study 2 piglets grouped by diagnosis (Healthy vs NEC). (A) Small intestinal contents. (B) Colon contents. Each stacked bar is one animal; colors denote species as indicated in the key, with low-abundance taxa collapsed into “Other”.

**Figure 10.**
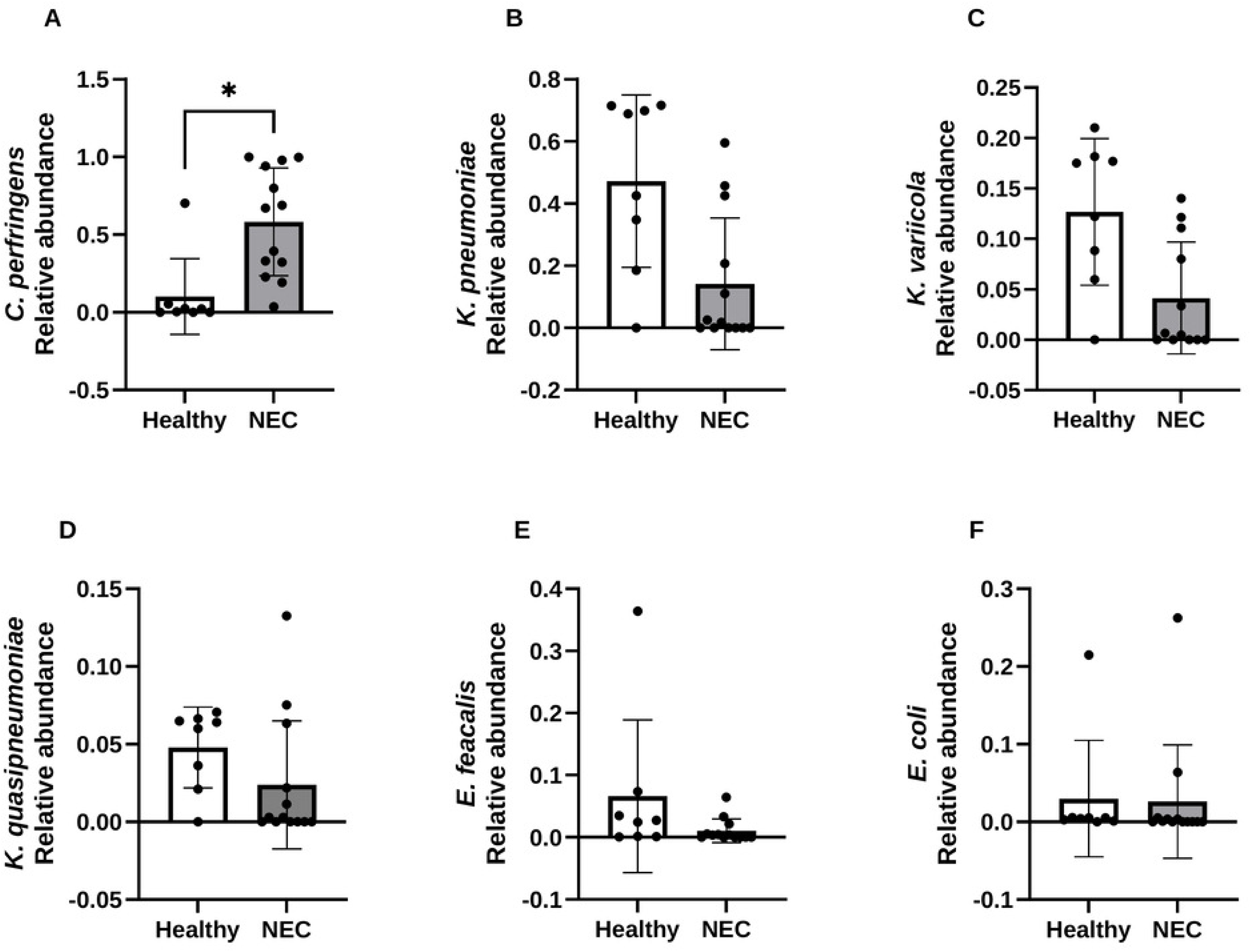
In the small intestine, NEC is associated with higher *Clostridium perfringens* and lower abundance of other taxa. Relative abundance of individual species in small intestinal contents of Study 2 piglets grouped by diagnosis (Healthy n = 8, NEC n = 12–13). (A) *Clostridium perfringens*. (B) *Klebsiella pneumoniae*. (C) *Klebsiella variicola*. (D) *Klebsiella quasipneumoniae*. (E) *Enterococcus faecalis*. (F) *Escherichia coli*. Each symbol is one animal and the bar is the group mean. Abundances were CLR-transformed and compared by Wilcoxon rank-sum test with Benjamini–Hochberg FDR correction (✱, P < 0.05).

**Figure 11.**
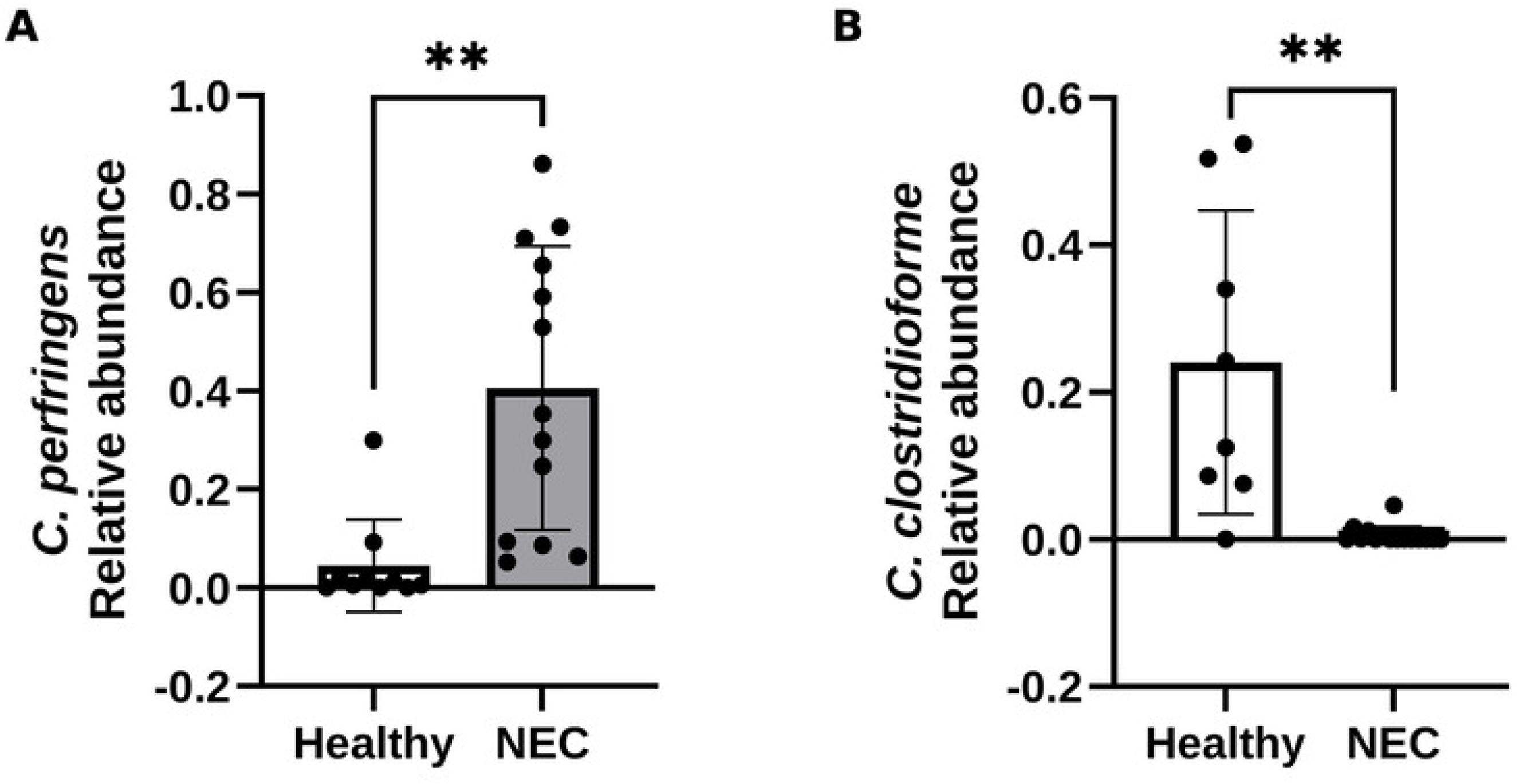
In the colon, NEC is associated with higher *Clostridium perfringens* and lower *Enterocloster clostridioformis*. Relative abundance of individual species in colon contents of Study 2 piglets grouped by diagnosis (Healthy vs NEC). (A) *Clostridium perfringens*. (B) *Clostridium clostridioforme* (*Enterocloster clostridioformis*). Each symbol is one animal and the bar is the group mean. Abundances were CLR-transformed and compared by Wilcoxon rank-sum test with Benjamini–Hochberg FDR correction (✱✱, P < 0.01).

## Discussion

The pathogenesis of NEC is incompletely understood, yet it is widely accepted that prematurity and microbial colonization of the intestine are two key risk factors for this disease. Gestational age is inversely correlated with the risk of developing NEC, with more premature infants at greater risk (11). Although NEC can be modeled in animals such as preterm piglets and mice, the disease does not develop in germ-free animals and is reduced with antibiotic treatment, providing strong evidence that the gut microbiota plays a critical role in its pathogenesis (12–14). Despite many years attempting to elucidate the relationship between NEC and the gut microbiota, there have been few robust associations established between specific bacterial species and disease incidence. However, one of the most consistent associations between specific taxa and the pathogenesis of NEC is an increase in Proteobacteria, namely Enterobacteriaceae, more commonly *Klebsiella* spp, preceding the onset of disease(2, 3, 15). Compounding this is evidence suggesting the involvement of toll-like receptor 4 (TLR4), which is recognizing Gram-negative-associated molecules like LPS and FimH, in the pathogenesis of NEC. TLR4 expression has been demonstrated to be higher in the intestinal tract of premature mice and infants compared to full term counterparts. This phenomenon is thought to be due to TLR4’s involvement in intestinal development, making this receptor more prevalent and active in preterm infants due to their underdeveloped intestinal epithelium at birth (4, 5). Moreover, studies in mice have demonstrated that knocking out TLR4 significantly decreases NEC-like intestinal injury in mouse pups subjected to a NEC induction protocol (5). Thus, in this study we set out to test whether oral treatment with sibofimloc, a FimH inhibitor, was safe and could ameliorate NEC incidence and severity in infant formula-fed preterm piglets. We hypothesized that blocking FimH adhesin, which allows efficient binding of gram-negative bacteria to epithelial cells, potentially eliciting inflammation in the vulnerable preterm intestine, would decrease colonization and further bloom of Enterobacteriaceae and reduce inflammatory events leading to the pathogenesis of NEC. To identify a safe and appropriate dose of sibofimloc for efficacy testing, we first conducted a pharmacokinetic and dose-escalation study in preterm piglets receiving 10, 25, or 50 mg/kg of Sibofimloc. Sibofimloc was absorbed into systemic circulation in a dose-dependent manner, yet its estimated fractional appearance remained low (<2% of the dose). This limited systemic bioavailability is intentional, as the therapeutic target for sibofimloc is localized within the gut lumen. Plasma sibofimloc concentrations declined between days 2 and 8 in all treatment groups, only significantly in the 50 mg/kg group, suggesting that age-dependent maturation of the intestinal epithelium and strengthening of the mucosal barrier reduce systemic absorption of the drug in the first week of life. Overall, despite the reduced enteral feeding volume, NEC incidence remained moderate (20–50%) across all sibofimloc dose groups. Although no overt adverse effects other than NEC were observed, it remains uncertain whether sibofimloc is safe in preterm piglets. More specific tolerance assays are necessary to determine the safety of sibofimloc in preterm piglets. We selected 40 mg/kg for the subsequent efficacy study to maximize intestinal drug exposure while avoiding the slight increase in NEC incidence observed in the 50 mg/kg group compared to the other doses.

Contrary to our hypothesis, oral treatment of formula-fed piglets with 40 mg/kg sibofimloc failed to reduce either NEC incidence or disease severity. Instead, sibofimloc-treated piglets exhibited a modest increase in NEC incidence (100% vs 75%) and more rapid NEC-associated mortality compared to CTL. This severity was evidenced by a modest decrease in villus height in sibofimloc compared to DHM and CTL groups and the significant decrease in crypt depth seen in sibofimloc-treated piglets. These histological findings in the intestinal tissue may indicate more severe destruction of the tissue that led not only to villus epithelial sloughing but also a potential loss of proliferative cells, and intestinal stem cells, impairing the regenerative capacity(16). Taken together this suggests that sibofimloc may worsen NEC disease severity in preterm piglets.

We evaluated the gut microbiota in small intestinal and colon contents when piglets either were diagnosed with NEC and euthanized or completed study without clinical signs of NEC. Whole genome sequencing (WGS) of contents from the small intestine and colon revealed that the gut microbiota of sibofimloc-treated piglets closely resembles that of CTL piglets but differed markedly from the microbiota in DHM-fed piglets. Although the difference in gut microbiota composition between CTL and DHM-fed piglets is a common finding in our studies (17). We found some gram-negative bacterial species in formula-fed piglets commonly observed in preterm infants (*Klebsiella sp. and E. coli*)(1, 2, 18). However, sibofimloc treatment did not significantly alter the gut microbiota composition compared to formula feeding alone. This may be expected given that the mechanism of action of sibofimloc is to prevent bacterial adherence and colonization on the epithelium. In striking contrast, piglets fed DHM exhibited a markedly higher abundance of *Klebsiella pneumoniae* (*K. pneumoniae*) compared to those fed formula. In our recent report (17), we also found that the DHM-fed piglets were heavily colonized with *Escherichia coli* (*E. coli*) regardless of NEC diagnosis compounding the theory that Enterobacteriaceae do not correlate with disease, but more so with health in this preterm piglet model. This is intriguing because this species is a member of the Enterobacteriaceae family and frequently associated with NEC disease(1, 2, 18). Slight differences in the relative abundance of *K. pneumoniae* between sibofimloc-treated and CTL piglets suggest that treatment may have inhibited colonization of this species, however, we cannot definitively conclude this from the data. Notably, the gut microbiota composition of these piglets suggests that regardless of treatment, there is a relatively low diversity of colonizing Enterobacteriaceae, mostly limited to *Klebsiella* species. Although this does not explain why sibofimloc did not reduce NEC, a possible reason is that our preterm piglet model does not exhibit Enterobacteriaceae-associated NEC.

Interestingly, in both sibofimloc and CTL groups, the dominant species within both the small intestinal and colonic microbiota was *Clostridium perfringens* (*C. perfringens*) which has previously been associated with NEC in the piglet model and multiple infant studies (12, 17, 19–23). The *C. perfringens* abundance was significantly higher in both CTL and sibofimloc-treated groups compared to DHM-fed piglets with a trend towards a higher average relative abundance of *C. perfringens* in sibofimloc-treated piglets. When comparing NEC-afflicted vs. healthy piglets, NEC was associated with higher abundance of *C. perfringens* and a less diverse microbial community. This is consistent with fact that *C. perfringens* does not express type 1 fimbriae, but sortase-assembled pilli and Type IV pilli, and thus sibofimloc should not inhibit their colonization (*24*). In contrast, healthy animals had more bacterial diversity, a modestly higher abundance of *Klebsiella spp* in the small intestine and a significantly higher abundance of *Clostridium clostridioforme*, recently reclassified as *Enterocloster clostridioformis* (*E. clostridioformis*) (25). Although we are unable to make robust conclusions about the significance of the association between a healthy outcome (no NEC) and *E. clostridioformis*, it is notable that this species is considered to be a commensal member of the microbiota with no reported virulence factors, unlike *C. perfringens*. In a recent study, *E. clostridioformis* monocolonization in germ-free mice was able to resist *Salmonella typhimurium* infection by promoting protective host responses within the intestinal epithelium and mucosa (26). These findings support the hypothesis that DHM supports the growth of microbial species (*Klebsiella spp* and *Enterocloster clostridioformis)* that may play a role in limiting NEC pathogenesis by preventing the expansion of *C. perfringens*.

Altogether, this study demonstrates that sibofimloc exhibits low systemic absorption and does not prevent NEC, but may exacerbate NEC in the preterm piglet model. Consistent with our previous studies, members of the Enterobacteriaceae were not strongly associated with disease in this preterm pig model, and sibofimloc treatment did not significantly alter the NEC-associated gut microbiota, which was predominantly characterized by an expansion of *C. perfringens*. These findings suggest that targeting FimH-expressing Enterobacteriaceae alone is insufficient to prevent NEC in this model, where disease appears to be driven primarily by non-FimH-producing pathogens. Importantly, this study provides the first pharmacokinetic and tolerability data for sibofimloc in a preterm animal model. The NEC incidence observed across all sibofimloc dose groups, despite reduced enteral feeding, indicates that this formulation or dosing strategy was not well tolerated under these experimental conditions. These findings highlight the importance of preclinical safety evaluation and will inform future optimization of sibofimloc for microbiota-associated diseases in preterm infants.

## Materials and Methods

### 1. Pharmacokinetics and Tolerability Assessment - Study 1

#### Animal Delivery, Clinical Care, Feeding Regimen

To assess the tolerability and pharmacokinetics of orally administered sibofimloc, 15 piglets were assigned to three treatment groups organized by escalating doses of oral SIBOFIMLOC given twice daily for 10 days after birth. The animal protocol was approved by the Institutional Animal Care and Use Committee (IACUC) of Baylor College of Medicine and was conducted in accordance with the National Institutes of Health (NIH) guidelines. Preterm piglets (n=16) were delivered at 103-104 days of gestation (term is 114 days) by cesarean section. A few hours after birth, the piglets were surgically implanted with a jugular venous catheter and an orogastric tube to facilitate parenteral nutrition and enteral feeding, respectively, as described previously (17, 27). Following surgery, piglets received total parenteral nutrition (TPN) via jugular vein at 50% of full nutritional requirements (120 mL/(kg*d), 410 kJ energy/(kg*d), 12.5 g dextrose/(kg*d), 6.5 g AA/(kg*d), 2.5 g fat/(kg*d). Orogastric feeding of the enteral formula (Enfamil Preterm 24) began on Day of Life (DOL) 2 at 60 ml/kg/d and continued until DOL 8 (Supplemental Table 1). The level of enteral feeding represents 25% of full feeding (240 ml/kg/d), done to minimize death from NEC during the pharmacokinetic and tolerability study given that disease is more commonly associated with higher levels of enteral feeding. Piglets were fed every 4 hours (6 times/day). The incidence of NEC monitored in all piglets. Piglets were monitored for overt adverse effects to sibofimloc including neurological events, respiratory distress, diarrhea, fever and weight loss.

##### Treatments

During the 8-day protocol, piglets were randomized into three treatment groups where sibofimloc was given via an orogastric feeding catheter at 10 mg/kg BID (Group 1), 25 mg/kg BID (Group 2), or 50 mg/kg twice daily (Group 3) (i.e. 20, 50, 100 mg/kg day). The sibofimloc was mixed in 0.5% (w/w) methylcellulose in deionized water. The sibofimloc doses began on DOL 1 and were given 12 h apart at 800 and 2000 hours each day and continued until DOL 10.

##### Pharmacokinetic Sampling Protocol

To quantify pharmacokinetics of sibofimloc, blood samples were collected on DOL 2 and 10 over a 12-h period post-dosing of compound among all study groups. On the morning of DOL 2 and 10, piglets were given their oral dose of sibofimloc. Blood samples (1 mL) were collected initially at 0 hours (prior to first dose) and again at 0.5, 1, 2, 4, 8, 12 hours post sibofimloc dosing. Plasma (∼500 uL) was obtained from whole blood samples and frozen (−80°C) for analysis of sibofimloc. Plasma concentrations of sibofimloc at DOL 2 and 10 were used to determine whether the compound was absorbed into systemic circulation. Blood collected from preterm piglets was used for sibofimloc assay development. To evaluate the proportion of drug absorption, the peak fraction of dose in blood was calculated for each animal and compared across levels and postnatal ages. For all animals, a blood volume of 70 ml/kg was assumed based on range reported for neonatal piglets (61-76 mg/kg) (28).

### 2. Efficacy of Sibofimloc to Prevent or Reduce NEC - Study 2 Animal Delivery, Clinical Care, Feeding Regimen

Based on the pharmacokinetics and tolerability study of sibofimloc (Study 1) a treatment dose of sibofimloc was established and tested for its efficacy in prevention of NEC. The animal protocol was approved by the Institutional Animal Care and Use Committee (IACUC) of Baylor College of Medicine and was conducted in accordance with the National Institutes of Health (NIH) guidelines. Preterm piglets (n=16) were delivered at 103-104 days of gestation (term is 114 days) by cesarean section. A few hours after birth, the piglets were surgically implanted with a jugular venous catheter and an orogastric tube to facilitate parenteral nutrition and enteral feeding, respectively as in Study 1. Following surgery, piglets received total parenteral nutrition (TPN) via jugular vein at 50% of full intake level (120 mL/(kg*d), 410 kJ energy/(kg*d), 12.5 g dextrose/(kg*d), 6.5 g AA/(kg*d), 2.5 g fat/(kg*d) (Intralipid) as previously described (17). Orogastric feeding of the enteral formula (Enfamil Preterm 24) began on Day of Life (DOL) 2 at 60 ml/kg/d (25% of full intake level) and increased gradually to 180 ml/kg/d (100% of full intake level) until DOL 11 (Supplemental Table 2). Piglets were fed every 4 hours (6 times/day) and monitored for symptoms of NEC.

#### Treatments

Preterm piglets (n=24) randomized into three treatment groups. For treatment with sibofimloc, a solution of 40mg/5mL of sibofimloc was prepared in phosphate-buffered saline (PBS) and a dose of 5mL/kg (equivalent to 40mg/kg) was administered to piglets every 12 hours in Enfamil Preterm 24. To ensure feeding regimen remained uniform, formula (CTL) and DHM groups received treatments of their respective diets and the PBS vehicle. Treatments began on DOL 2 enterally via orogastric catheters and continued until DOL 11.

### 3. NEC Monitoring

Piglets received regular, standardized physical exams to evaluate for clinical symptoms of NEC including abdominal distension, lethargy, cyanosis and bloody diarrhea. Piglets that developed signs of NEC were monitored by SpO2 and when readings dropped below 80% for more than 3-4 hours, piglets were euthanized as described previously (17, 29).

### 4. Tissue and Intestinal Content Sample Collection

On DOL 11, all remaining piglets were euthanized, piglets in sibofimloc group were euthanized 4 hours after their morning dose of sibofimloc. Plasma samples were collected before euthanasia. Upon euthanasia, samples of small intestinal and colon contents were collected. All tissue, plasma and content samples were stored at -80°C immediately upon collection. Formalin fixed and OCT frozen tissues were collected from distal ileum and colon. Gastrointestinal tract segments (stomach, proximal jejunum, distal ileum and colon) were scored macroscopically using a standardized scoring system (29) resulting in a gross NEC score. Formalin-fixed samples were used for histopathological analysis using H&E staining was performed using Zeiss Blue version 3.3, resulting in a histological NEC score using a system described previously (17).

### 5. Microbiota Analysis

Metagenomic whole genome shotgun (WGS) sequencing of final small intestinal and colon contents was completed by the Baylor College of Medicine Alkek CMMR. Metagenomic whole genome shotgun (WGS) sequencing allows for species-level resolution of taxa and functional enzyme/metabolic pathway content. Briefly, genomic DNA is extracted using Qiagen DNeasy Power Soil Pro HT. Libraries are constructed using Illumina DN Prep. Completed libraries undergo quality control using a combination of PicoGreen, Qubit, Fragment Analyzer and Tapestation to assess concentration and fragment size distribution. Shotgun sequencing of pooled libraries is carried out using Illumina NovaSeq platform using 2x150bp paired-end protocol. Demultiplexed raw FASTQ sequences are processed using BBDuk and trimmed FASTQs are mapped to a combined PhiX and host reference genome database using BBMap. Processed sequences are taxonomically profiled using MetaPhlAn3. Functional gene and metabolic pathway profiles are generated using HUMAnN2 (30).

### 6. Statistical Analysis

Results were analyzed using GraphPad Prism 10. The following statistical analyses were performed to determine significant differences for a given observation: For all comparisons of NEC incidence a Fisher’s exact test was used to determine the difference between groups; one-way ANOVA with Tukey’s multiple comparison was performed for comparison of gross and histological NEC severity scores; villus height and crypt depth measurements were compared using unpaired t test with Welch’s correction for comparisons between diagnosis and one-way ANOVA with Tukey’s multiple comparisons test was performed to compare groups; area under the curve (AUC) values of doses between day 2 and day 8 were compared using Mann-Whitney tests while comparisons in AUC between doses were performed using a one-way ANOVA. Relative abundance data were transformed using a centered log-ratio (CLR) transformation to account for the compositional nature of microbiota sequencing data. CLR-transformed abundances were calculated in R (v4.4.2) using the compositions package. Statistical analyses were performed on CLR-transformed abundances. Differences in bacterial abundances among treatment groups were assessed using the Kruskal–Wallis test, followed by pairwise Wilcoxon rank-sum tests when appropriate. Comparisons between NEC and healthy animals were performed using the Wilcoxon rank-sum test. To account for multiple comparisons across taxa, *P* values were adjusted using the Benjamini–Hochberg false discovery rate (FDR) correction, with an adjusted *P* < 0.05 considered statistically significant. Peak fraction in blood (%) comparisons were performed by log-transformation of percentages followed by two-way ANOVA with Šídák’s multiple comparisons test.

## Acknowledgments

The authors would like to acknowledge Inka Didelija, Liwei Cui, and Xiaoyan Chang for their hard work in completing laboratory assays and animal care, which contributed significantly to the completion of this project.

## Author contributions

**VMH:** Methodology, Investigation, Formal analysis, Data curation, Visualization, Writing and Editing

**DB:** Conceptualization, Methodology, Supervision, Funding acquisition, Writing, Reviewing and Editing

**BS:** Conceptualization and Methodology

**BE:** Investigation, and Writing

**HH, SKB, MP and MH:** Conceptualization, Methodology, Formal analysis, Reviewing

**GG:** Investigation and Methodology

**CV:** Investigation, Reviewing and Editing

## Supplemental Material

1. Supplemental Table 1. Piglet parenteral and enteral nutrition for pharmacokinetic study (Study 1)

2. Supplemental Table 2. Piglet parenteral and enteral nutrition for efficacy study (Study 2)

